# Dietary α-Ketoglutarate delays lung adenocarcinoma in females and modulates TBX5-associated transcriptional programs and the immune microenvironment

**DOI:** 10.1101/2025.09.19.677388

**Authors:** Martín Alcaraz, Aparamita Pandey, Mia Santiago, Camelia Dumitras, Leilani Pradis, Estefany Gomez, Adriana Soto, Pasquale Saggese, Bin Liu, Steven M. Dubinett, Claudio Scafoglio

## Abstract

Lung adenocarcinoma (LUAD) is the most common form of lung cancer and a leading cause of cancer-related mortality, underscoring the need for new chemopreventive strategies. α-Ketoglutarate (α-KG), a tricarboxylic acid cycle metabolite and dioxygenase cofactor, links cellular metabolism to chromatin regulation. Here, we show that dietary α-KG remodels LUAD in a sex-dependent manner. In female mice, α-KG reduced tumor area, decreased repressive histone marks (H3K27me3, H3K9me3), and upregulated TBX5 and myogenesis-associated genes. In male mice, α-KG-treated male mice exhibited increased tumor area, elevated H3K27me3, and immune remodeling characterized by CD8⁺ T cell expansion and transcriptomic signatures of T cell exhaustion. Analysis of human LUAD revealed that TBX5 expression is enriched in female tumors and associated with improved survival, suggesting it may serve as a marker of favorable outcome. Together, these findings support α-KG as an epigenetic modulator with potential chemopreventive activity in lung cancer and highlight the importance of incorporating sex as a biological variable in preclinical studies.

**Summary:** Oral α-Ketoglutarate (α-KG) has a known anti-aging effect and has been suggested to inhibit cancer development. However, the role of α-KG in lung cancer is not known. Here, we investigated the effect of oral α-KG on the development of lung adenocarcinoma in a Kras*^G12D^*-driven murine model. In females, α-KG reduced lung tumor growth, accompanied by TBX5 induction, activation of a myogenesis transcriptional program, and loss of repressive histone methylation. In males, α-KG increased tumor growth, coinciding with TBX5 repression, suppression of myogenesis programs, and accumulation of repressive histone methylation. α-Ketoglutarate also remodeled the tumor immune microenvironment in a sex-dependent manner, with divergent effects on T cell infiltration.

**HIGHLIGHTS:**

- Oral α-Ketoglutarate reduces lung adenocarcinoma development in a genetically engineered murine model in a sex-dependent manner
- In females, α-Ketoglutarate reduces repressive histone marks and induces TBX5/myogenesis
- In males, α-Ketoglutarate increases repressive histone marks and suppresses TBX5/myogenesis
- α-Ketoglutarate remodels the tumor immune microenvironment with sex-specific effects
- TBX5 is enriched in female LUAD patients and predicts improved survival in human datasets

**CONTEXT AND SIGNIFICANCE:** Lung adenocarcinoma (LUAD) is the most common form of lung cancer and a leading cause of cancer mortality worldwide. Preventive strategies are limited, highlighting the need for safe approaches that can intercept tumor progression. Metabolic cofactors are increasingly recognized as modulators of the cancer epigenome. Among these, α-Ketoglutarate (α-KG), a central metabolite of the tricarboxylic acid cycle, functions as a cofactor for chromatin-modifying enzymes. Here, we demonstrate that dietary α-KG supplementation limits LUAD progression in female mice, reduces repressive histone methylation, and activates TBX5-associated transcriptional programs. These effects are sex dependent, underscoring the importance of biological context in shaping metabolic responses. In human LUAD, TBX5 is enriched in females and predicts improved survival, suggesting that it may serve as a marker of favorable outcome and help guide future studies of α-KG. Together, this work identifies α-KG as an epigenetically active metabolite with translational promise for lung cancer chemoprevention.

**GRAPHICAL ABSTRACT:** 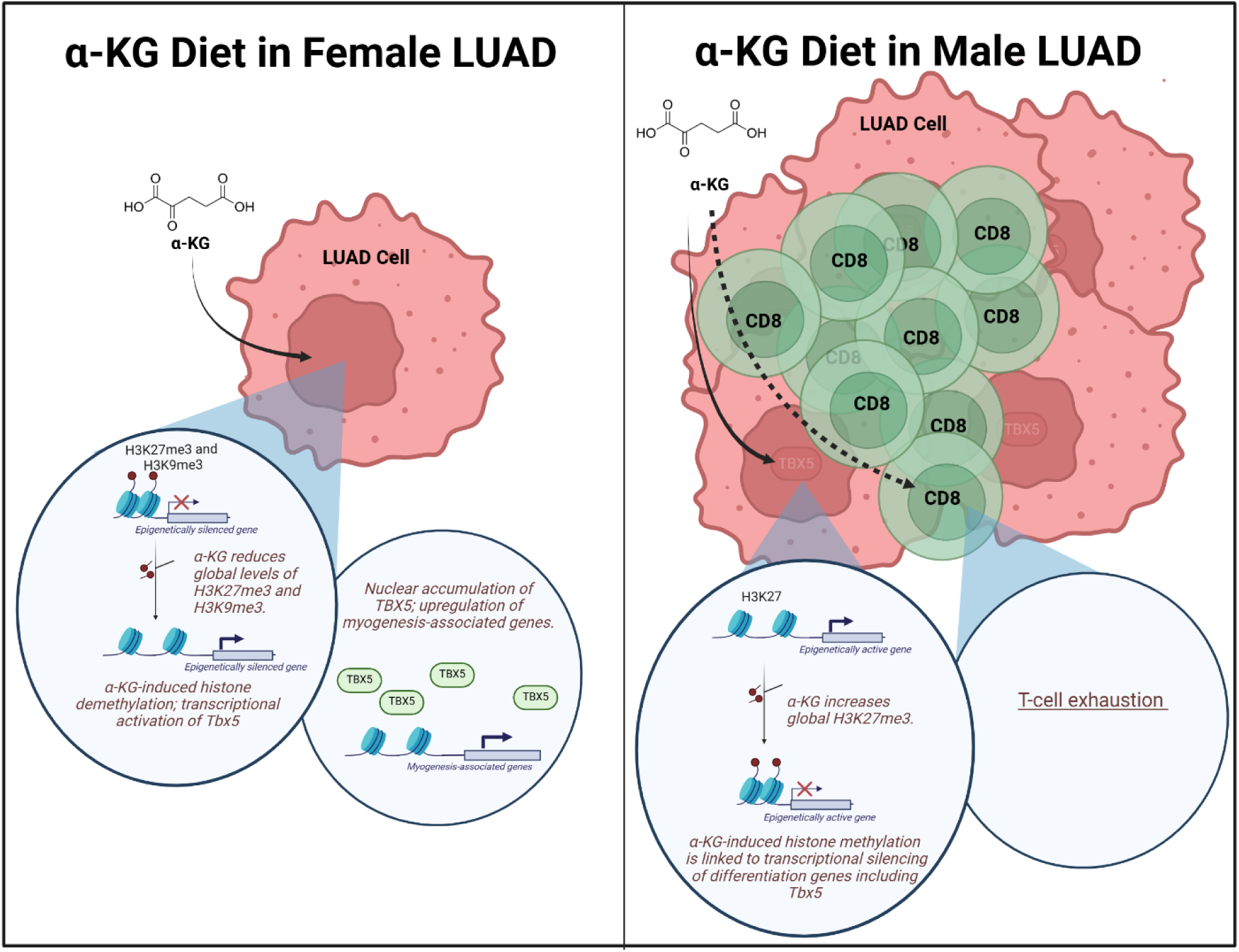

**One-sentence summary:** α-Ketoglutarate remodels lung adenocarcinoma through sex-dependent epigenetic, transcriptional, and immune reprogramming.

## INTRODUCTION

Lung cancer remains the leading cause of cancer-related death worldwide, with five-year survival rates still poor despite advances in targeted therapies and immunotherapies^1^. Lung adenocarcinoma (LUAD) is the most frequent type of lung cancer. Activating mutations in *KRAS*, particularly the G12D allele, are among the most common oncogenic drivers in LUAD^2^. While tumor-intrinsic mutations are critical, LUAD progression is also shaped by interactions between cancer cells, their metabolic environment, and the host immune response^3–5^. Metabolic interventions, including dietary and pharmacologic approaches, are now being tested in clinical trials across multiple cancers^6–8^. Understanding how metabolites influence the tumor epigenetic state and progression may therefore have direct translational relevance.

α-Ketoglutarate (α-KG), a central metabolite in the tricarboxylic acid (TCA) cycle, is a required cofactor for α-KG-dependent dioxygenases that regulate diverse cellular processes. These include histone, DNA, and RNA demethylation, as well as other enzymatic activities that link cellular metabolism to gene regulation^9–14^. Prior studies have shown that α-KG can enhance stem cell pluripotency, modulate immune responses, and influence cancer cell plasticity^15–17^. However, the in vivo impact of α-KG on tumor progression, particularly in *KRAS*-driven LUAD, remains largely unexplored.

Here, we investigated the effects of dietary α-KG supplementation on tumor biology in a Kras*^G12D^*-driven genetically engineered mouse model of LUAD. Our findings reveal sex-dependent differences in tumor growth, histone methylation, and immune remodeling, and highlight transcriptional changes associated with TBX5 and myogenesis-related gene expression in females. These results position α-KG as an epigenetically active compound with potential chemopreventive relevance in LUAD, while underscoring that sex and tumor genotype are critical biological variables that must be incorporated into preclinical studies.

## RESULTS

### Dietary α-Ketoglutarate Reduces Tumor Area in Female LUAD Mice but Not in Males

To test whether α-ketoglutarate (α-KG) supplementation influences lung adenocarcinoma (LUAD) progression, we used a conditional genetically engineered murine model driven by *Kras^LSL-G12D/+^* on C57BL/6J background. Tumors were initiated via intranasal delivery of adenovirus expressing Cre recombinase (AdCre) at 12 weeks of age, and mice were randomized one week later to receive either a control diet or a diet supplemented with 2% calcium α-KG (Ca-αKG) for 16 weeks (Fig. 1A).

**Figure 1.**
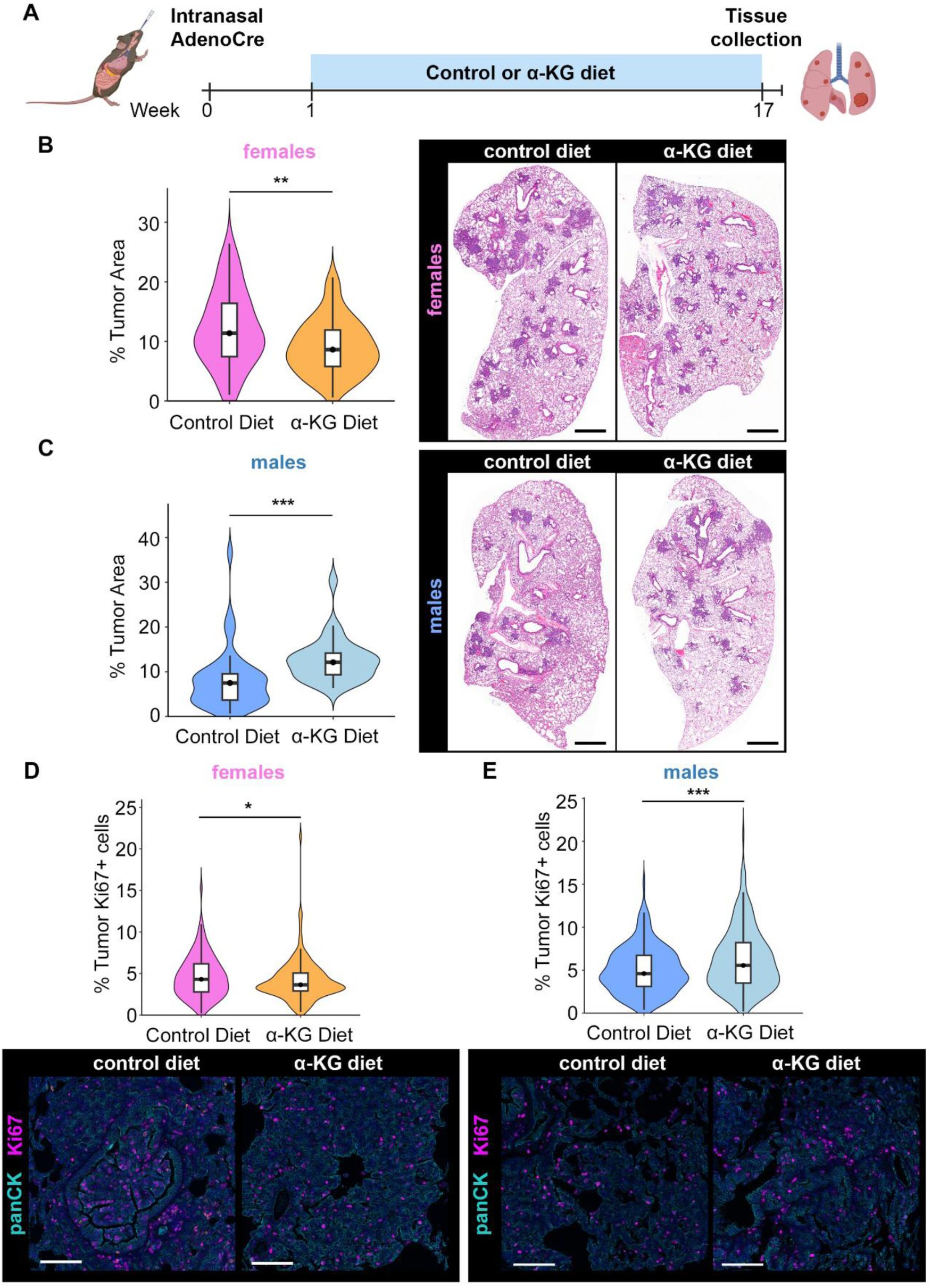
Dietary α-Ketoglutarate Reduces LUAD Tumor Area in Female but Not in Male Mice. **(A)** Schematic of the in vivo dietary intervention experiment. Mice were administered intranasal AdenoCre at 12 weeks of age to initiate lung adenocarcinoma and placed on either a control diet or 2% (w/w) calcium α-ketoglutarate (Ca-αKG) diet beginning at week 1 post-infection (n = 13 female and 10 male mice for control diet, 16 female and 10 male mice for α-KG). Lungs were harvested at week 17 for analysis. **(B-C)** Representative H&E-stained lung sections from female (B) and male (C) mice on control or Ca-αKG diet (right), with quantification of tumor area (left). Significance was evaluated by unpaired Student’s *t*-test (B) and Wilcoxon rank-sum test (C). Scale bars = 1 mm. **(D–E)** Quantification of proliferating tumor cells (top) and representative multiplex immunofluorescence (mIF) images (bottom) of PanCK^+^ (cyan) Ki67^+^ (magenta) tumor cells in female (D) and male (E) LUAD tumors. Tumor proliferation was assessed as % PanCK^+^ Ki67^+^ cells by multiplex immunofluorescence. Significance was estimated by Wilcoxon rank-sum test. Scale bars = 100 μm.

Tumor area was quantified from hematoxylin and eosin (H&E)-stained lung sections. At baseline, female LUAD tumors were significantly larger than male tumors (p = 0.0002; Fig. S1). In female mice, Ca-αKG significantly reduced tumor area (p = 0.0025; Fig. 1B), whereas in male mice, Ca-αKG increased tumor area compared to the control diet (p < 0.0001; Fig. 1C). Analysis of tumor cell proliferation by multiplex immunofluorescence (mIF) showed that Ca-αKG reduced proliferation in females (p = 0.05; Fig. 1D) but significantly increased proliferation in males (p = 0.0025; Fig. 1E).

These results demonstrate a sex-dependent effect of dietary α-KG on LUAD, with opposing outcomes in females versus males.

### Dietary α-Ketoglutarate Induces a Myogenesis Transcriptional Program in Female Tumors

To further investigate the effect of dietary αKG on lung cancer, we performed bulk RNA sequencing of LUAD tumors from mice on control or Ca-αKG diet. This dataset first allowed us to examine baseline sex-dependent transcriptomic differences in tumors from mice on a control diet. Gene set enrichment analysis (GSEA) identified multiple pathways with sex-biased activity, including inflammatory and interferon responses elevated in females and myogenesis, oxidative phosphorylation, and *Myc* signaling enriched in males (Fig. S2). These findings suggest distinct immune-metabolic states that may influence tumor responses to α-KG.

We next compared α-KG vs control diet within each sex. Strikingly, the only pathway that reached significance was the myogenesis gene set, which was strongly enriched in α-KG-treated female tumors (NES = 1.57; FDR q < 0.001; ∼20% leading-edge contribution; Fig. 2A). In contrast, the same pathway was significantly suppressed in males (NES = –0.93; FDR q < 0.001; ∼19% leading edge; Fig. 2B). A heatmap of myogenesis-related transcripts highlighted induction of *Mybpc3*, *Actn2*, *Hrc*, *Tnnt2*, *Tnnc1*, and *Myom1* in females (Fig. 2C). Consistently, volcano plots identified sex-dimorphic upregulation of *Myom2*, *Hrc*, *Fabp3*, *Ldb3*, and *Tnnt1* in females (Fig. 2D). These genes are typically restricted to cardiac or skeletal muscle and are rarely expressed in epithelial tumors ^18–20^. Complete differential gene expression results are provided in Supplementary Table S1.

**Figure 2.**
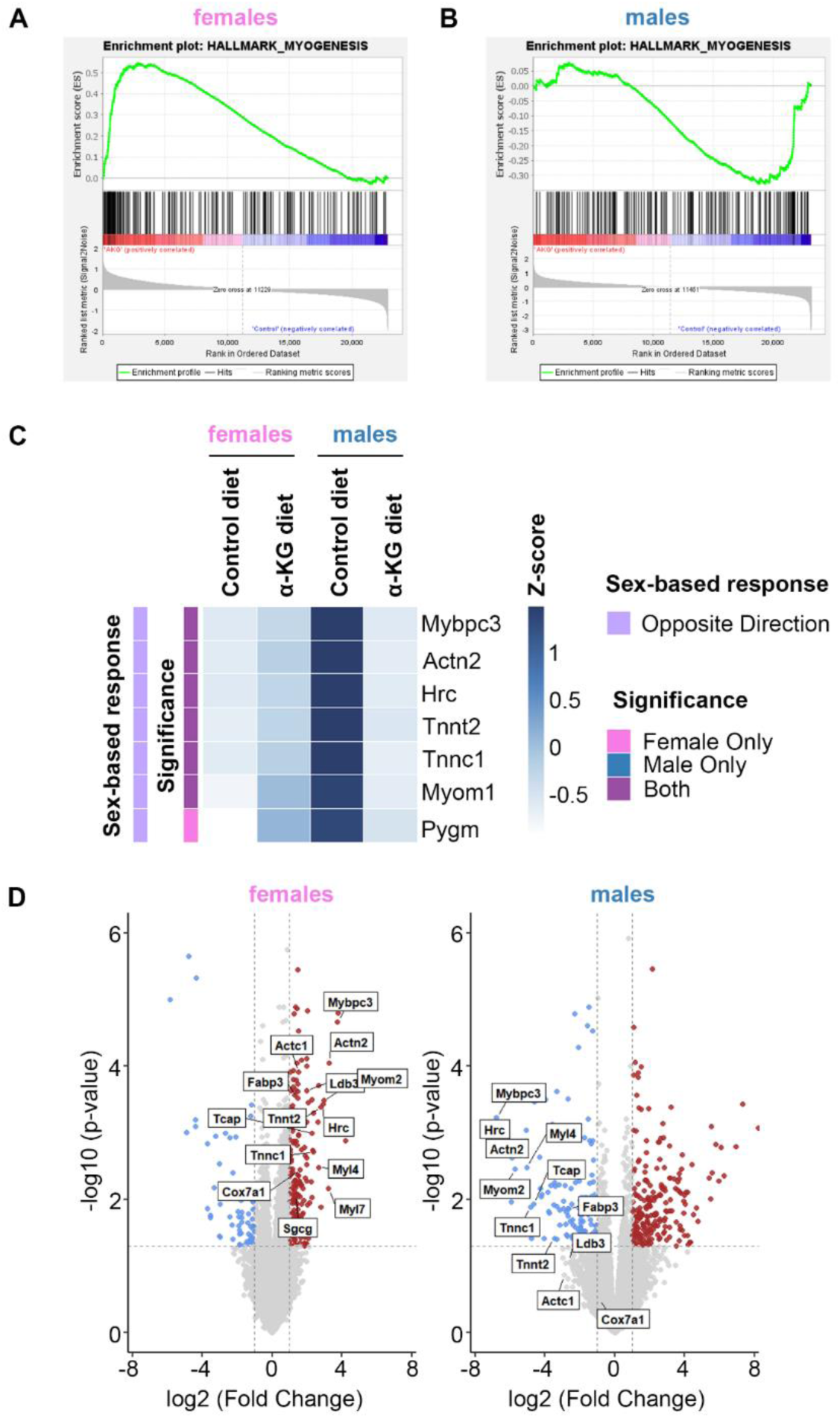
Dietary α-Ketoglutarate Induces a MYOGENESIS Transcriptional Program in Female Lung Tumors. **(A-B)** Gene Set Enrichment Analysis (GSEA) of the HALLMARK_MYOGENESIS gene set in female (A) and male (B) tumors comparing Ca-αKG diet vs control diet. **(C)** Heatmap of selected MYOGENESIS-related genes across sexes and diets. Genes are annotated by significance (female-only, male-only, or both) and direction of sex-based response. **(D)** Volcano plots of differential gene expression (Ca-αKG vs control diet) for female (left) and male (right) LUAD tumors. Genes are colored based on statistical significance (red: up-regulated; blue: down-regulated). Myogenesis genes are labelled. Data are from bulk RNA sequencing of lung tumors harvested at 17 weeks (n = 4 tumors per group). Differential expression was performed using DESeq2. GSEA was performed using the Hallmark gene set collection (MSigDB v7.5.1). NES = normalized enrichment score. Significance cutoff: adjusted *p*-value < 0.05.

Together, these results show that dietary α-KG uniquely induces a myogenesis-related transcriptional program in female LUAD tumors, whereas the same pathway is suppressed in males.

### TBX5 Is Associated with a Myogenesis Transcriptional Program in α-KG-Treated Female LUAD Tumors

To identify upstream regulators of α-KG-associated transcriptional changes, we performed ChEA3 transcription factor enrichment analysis ^21^ using leading-edge myogenesis genes. TBX5 and GATA4 emerged as top candidate regulators. The full ChEA3 transcription factor enrichment output is available in Supplementary Table S2.

Immunohistochemistry (IHC) for TBX5 showed increased nuclear staining in α-KG-treated female tumors (p = 0.0009; Fig. 3A) and decreased expression in males (p = 0.0004; Fig. 3B). GATA4 levels were unchanged in females (p = 0.218; Fig. 3C) but significantly increased in males (p = 4.65e-08; Fig. 3D), indicating a sex-specific pattern of transcription factor regulation.

**Figure 3.**
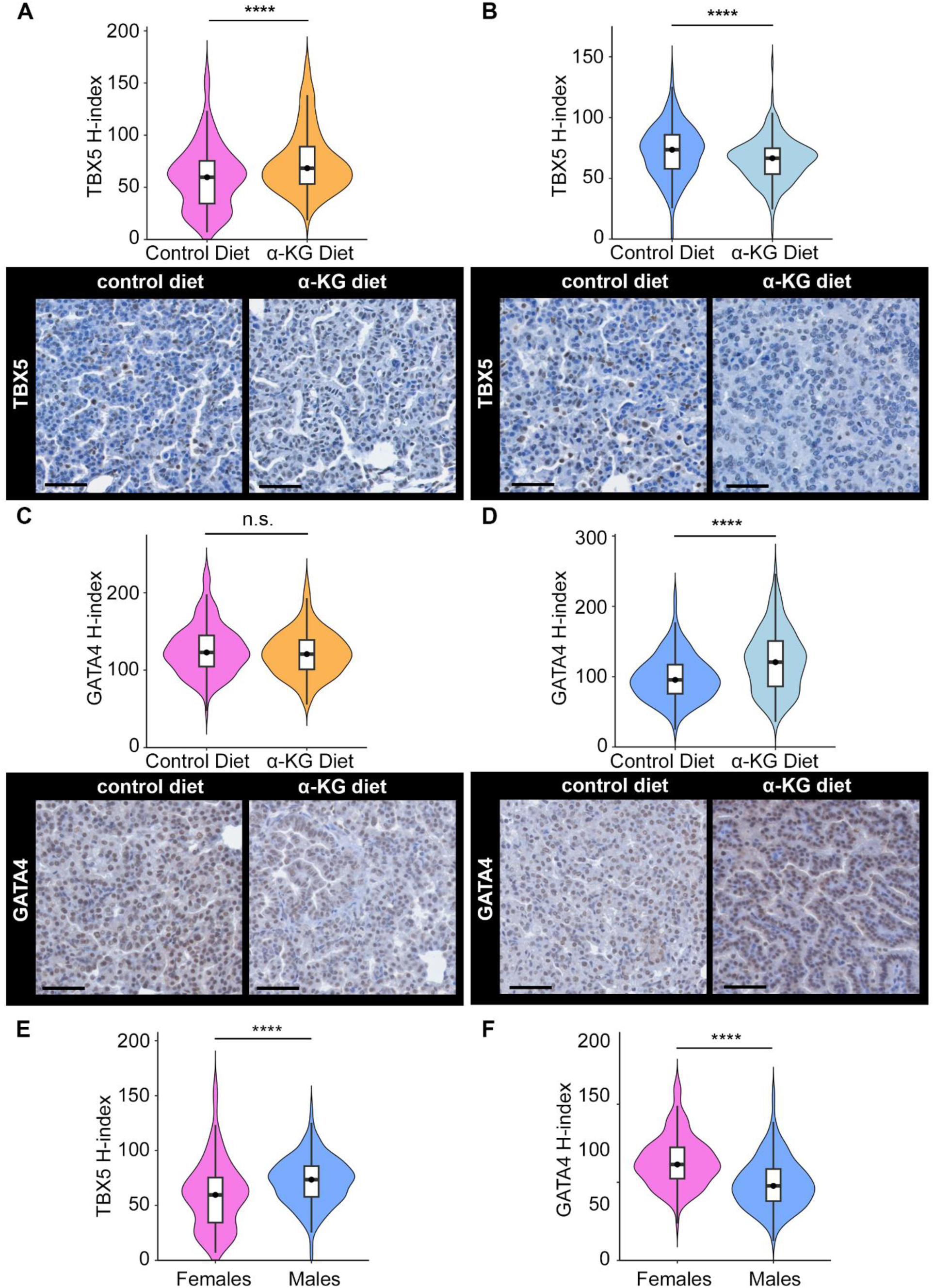
TBX5 Is Associated with a MYOGENESIS Transcriptional Program in α-KG-Treated Female LUAD Tumors. **(A-B)** Quantification of TBX5 nuclear expression by IHC (H-score) in LUAD tumors (n = 4 whole-lung sections per group) from (A) female and (B) male mice. Representative IHC images are shown below. Scale bars = 50 µm. **(C-D)** Quantification of GATA4 nuclear expression by IHC (H-score) in LUAD tumors (n = 4 whole-lung sections per group) from (C) female and (D) male mice. Representative IHC images are shown below. Scale bars = 50 µm. **(E–F)** Quantification of baseline TBX5 (E) and GATA4 (F) expression by IHC (H-score) in LUAD tumors from female and male mice on normal diet.

To contextualize these findings, we next compared baseline TBX5 and GATA4 expression between sexes on normal diet (Fig. 3E–F). Baseline TBX5 expression was significantly higher in males (p = 5.16e-5), whereas GATA4 expression was significantly higher in females (p = 6.70e-13).

Together, these results identify TBX5 as a sex-biased transcription factor associated with α-KG treatment in female LUAD tumors. GATA4 showed less consistent changes, but baseline differences in both transcription factors indicate that sex influences their relative abundance. These baseline disparities may shape transcriptional responses to α-KG and help explain activation of the myogenesis program in females.

### Dietary α-Ketoglutarate Alters Repressive Histone Methylation in a Sex-Dependent Manner

To assess changes in global histone methylation patterns, we examined H3K27me3, H3K9me3, and H3K4me3, which are histone modifications previously shown to be sensitive to α-KG availability and linked to LUAD dedifferentiation and aggressiveness^22^. Western blotting of histone extracts from LUAD tumors revealed that in females, α-KG supplementation significantly reduced H3K27me3 (p = 0.05) and H3K9me3 (p = 0.04), while H3K4me3 showed a downward trend that did not reach significance (Fig. 4A-B). In males, α-KG treatment significantly increased H3K27me3 (p = 0.002), with no change in H3K4me3 or H3K9me3 (Fig. 4C-D).

**Figure 4.**
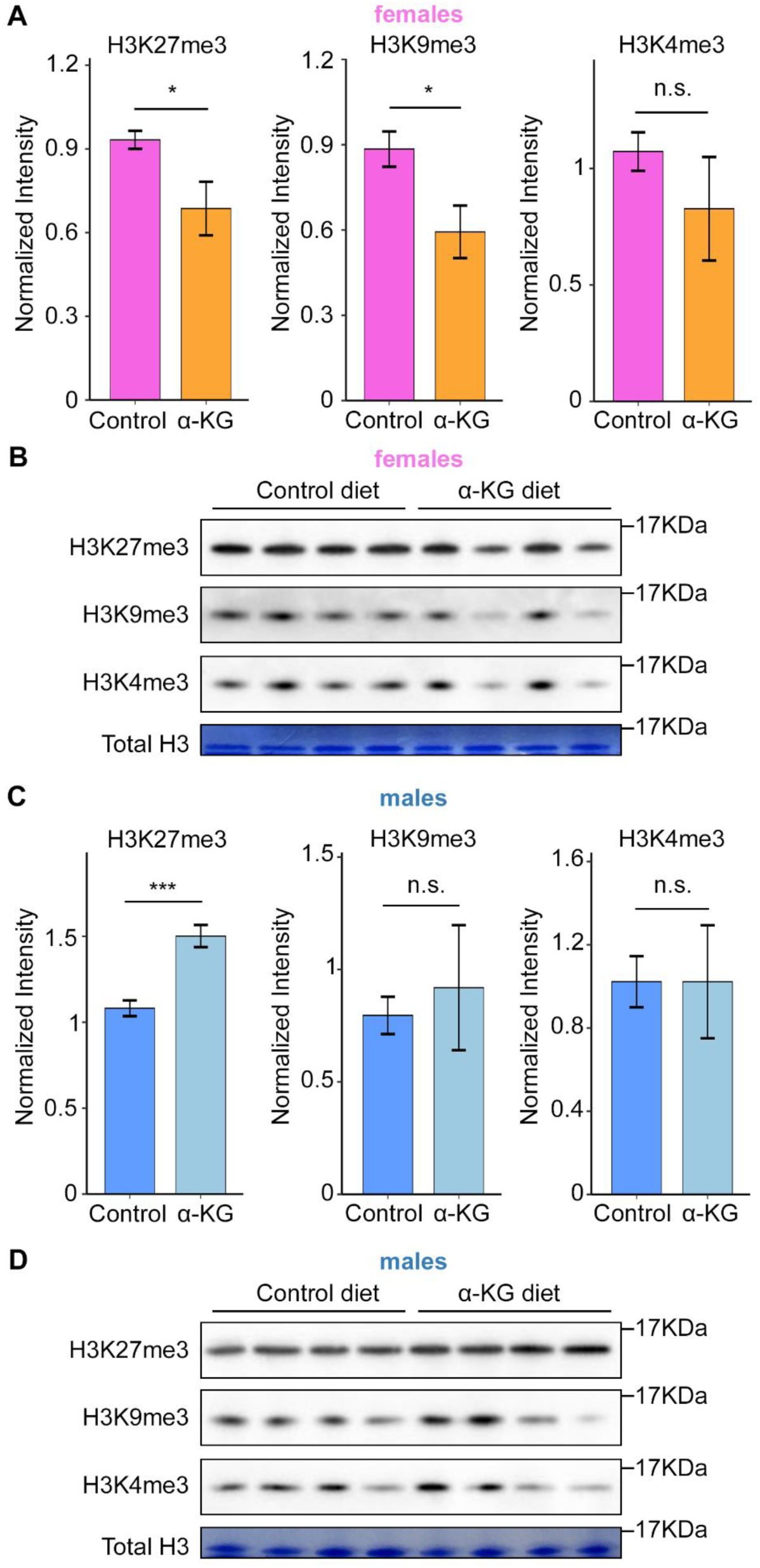
Dietary α-Ketoglutarate Alters Repressive Histone Methylation in a Sex-Dependent Manner. **(A–B)** Quantification (A) and representative western blot (B) of histone methylation levels (H3K9me3, H3K4me3, H3K27me3) in lung tumors from female mice on control or Ca-αKG diet. **(C–D)** Quantification (C) and representative western blot (D) of histone methylation levels in lung tumors from male mice on control or Ca-αKG diet. Total histone input was visualized by Coomassie staining at the corresponding molecular weight range and used for normalization. Panels are shown separately for clarity. Data are presented as mean ± SEM. n = 4 mice per group. Unpaired *t*-tests were used for all comparisons.

These results indicate that dietary α-KG alters repressive histone methylation in a sex-dependent manner, reducing repressive marks in females while increasing them in males. Each Western blot lane represents an independent tumor sample (n = 4 mice per group), and the concordance of these findings with transcriptional and IHC data supports a consistent biological effect.

### TBX5 Expression Associates with Lineage Programs, Proliferation, and Prognosis in Human LUAD

To extend our findings to human disease, we analyzed TBX5 expression in The Cancer Genome Atlas (TCGA) LUAD dataset. *TBX5* levels were significantly higher in female tumors compared to male tumors (p = 0.035, Fig. S3A). Notably, this sex bias differs from mouse baseline tumors, underscoring the context-dependent nature of TBX5 regulation. Kaplan-Meier survival analysis showed that high *TBX5* expression was associated with improved five-year overall survival in female patients (p = 0.028) and exhibited a similar but non-significant trend in males (p = 0.28) (Fig. 5A-B). Cox proportional hazards modeling confirmed that low *TBX5* expression predicted worse survival in females (HR = 1.58, p = 0.03), while in males, the similar trend did not reach statistical significance (HR = 1.28, p = 0.30) (Fig. 5C-D).

**Figure 5.**
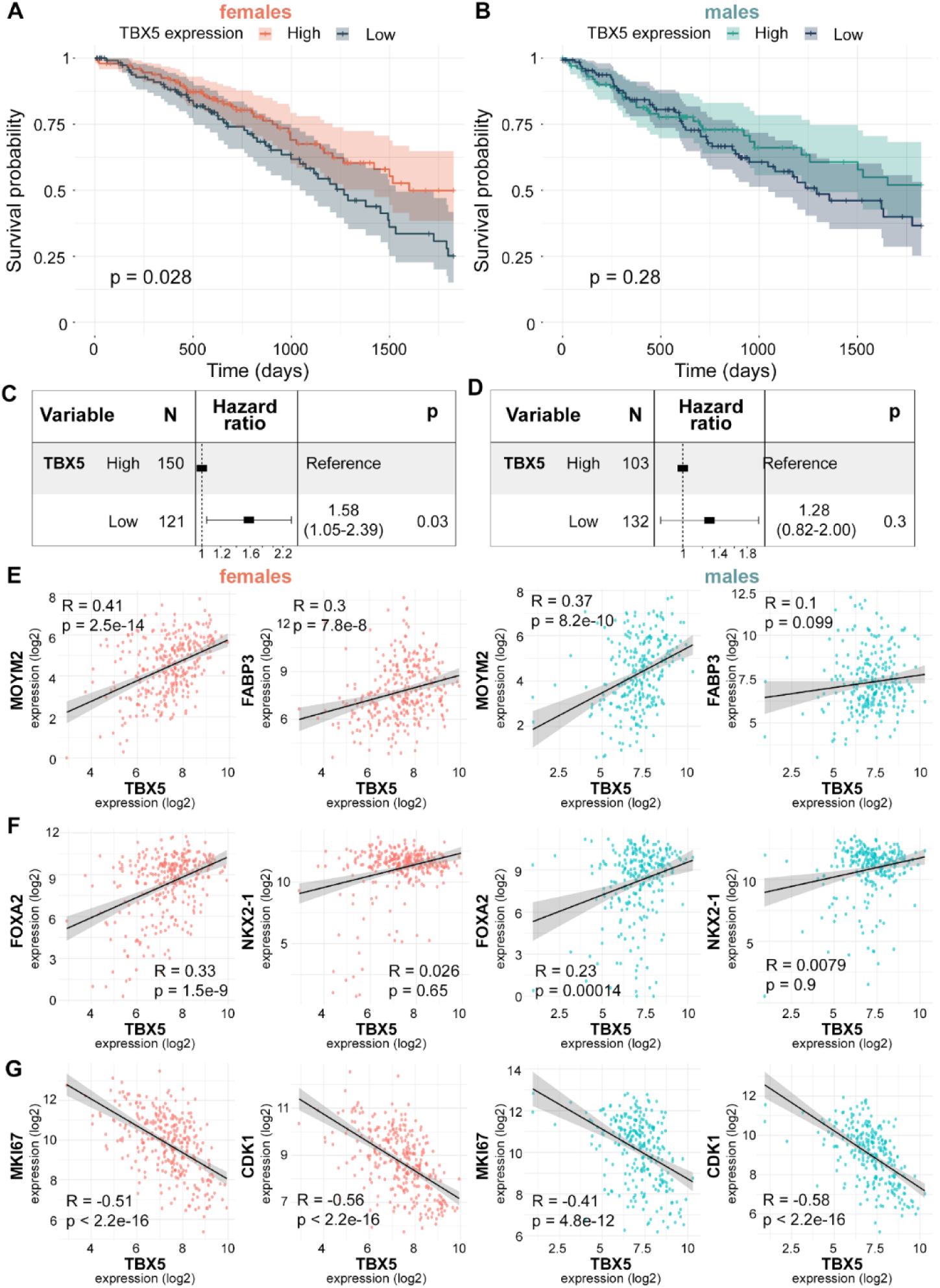
TBX5 Expression Associates with Lineage Programs, Proliferation, and Prognosis in Human LUAD. **(A-B)** Kaplan–Meier survival curves stratified by TBX5 expression (median split) in females (log-rank *p* = 0.028) and in males (log-rank *p* = 0.28). **(C-D)** Cox proportional hazards modeling of TBX5 correlation with 5-year survival in females (HR = 1.58, 95% CI: 1.05–2.39, *p* = 0.03) and in males (HR = 1.28, 95% CI: 0.82–2.00, *p* = 0.3). **(E-G)** Correlation of TBX5 expression with myogenesis genes (E), with markers of well-differentiated LUAD (F), and with proliferation markers (G) in female (left panels) and male (right panels) LUAD tumors. Data were obtained from TCGA LUAD RNA-seq and clinical annotations via UCSC Xena. *P*-values were calculated using two-sided Wilcoxon rank-sum tests, log-rank tests, or Cox proportional hazards models as indicated. Correlations represent Spearman’s rank coefficients with fitted trend lines and 95% confidence intervals.

To further examine the biological context of *TBX5* in LUAD, we assessed transcriptional correlations using TCGA data. *TBX5* expression was positively correlated with myogenesis genes *MYOM2* (R = 0.41 in females, p = 2.5e–14; R = 0.37 in males, p = 8.2e–10) and *FABP3* (R = 0.30 in females, p = 7.8e–08; R = 0.10 in males, p = 0.099), as well as the marker of well-differentiated LUAD, *FOXA2* (R = 0.33 in females, p = 1.5e–09; R = 0.23 in males, p = 0.00014). Correlation with the LUAD marker *NKX2-1* was positive but weak and not statistically significant (R = 0.026 in females, p = 0.65; R = 0.0079 in males, p = 0.90). *TBX5* expression was strongly and inversely correlated with proliferation markers *MKI67* (R = – 0.51 in females, p < 2.2e–16; R = –0.41 in males, p = 4.8e–12) and *CDK1* (R = –0.56 in females, p < 2.2e–16; R = –0.53 in males, p < 2.2e–16) (Fig. 5E–G).

Together, these findings suggest that *TBX5* is associated with a more lineage-committed and less proliferative tumor state. Clinically, high *TBX5* expression was significantly associated with improved survival in female patients and showed a similar, though non-significant, trend in males. In contrast, *GATA4* expression was higher in female LUAD tumors and high *GATA4* expression was associated with worse 5-year survival in female patients (p = 0.046). This effect was not observed in males (p = 0.41), indicating that GATA4 does not provide consistent prognostic information across sexes (Fig. S3B-D).

### α-Ketoglutarate Remodels the Tumor Immune Microenvironment in a Sex-Dependent Manner

To evaluate immune cell changes, we first performed IHC for CD3^+^ T cells (pan-T cell marker recognizing both CD4^+^ and CD8^+^ subsets). Dietary α-KG significantly reduced CD3^+^ infiltration both in females (p = 0.0005; Fig. 6A) and in males (p = 0.002; Fig. 6B). To further characterize the T cell populations in the immune microenvironment, we performed multiplex immunofluorescence (mIF). In males, α-KG significantly increased CD8^+^ T cells, whereas they were unchanged in females. CD4^+^ T cells were very rare (less than 0.5% of cells in the tumor microenvironment) and were significantly decreased in both sexes (Fig. 6C-D). A reduction in CD3^+^ T cells in both sexes and increase in CD8+ T cells in males appeared inconsistent with the observed tumor burden changes. However, analysis of the bulk RNA-seq data showed down-regulation of multiple immune checkpoint genes (*Pdcd1*, *Ctla4*, *Lag3*) more pronouncedly in females and up-regulation of checkpoint gene *Tigit* in males (Fig. S4A), suggesting functional changes in the infiltrating T cells.

**Figure 6.**
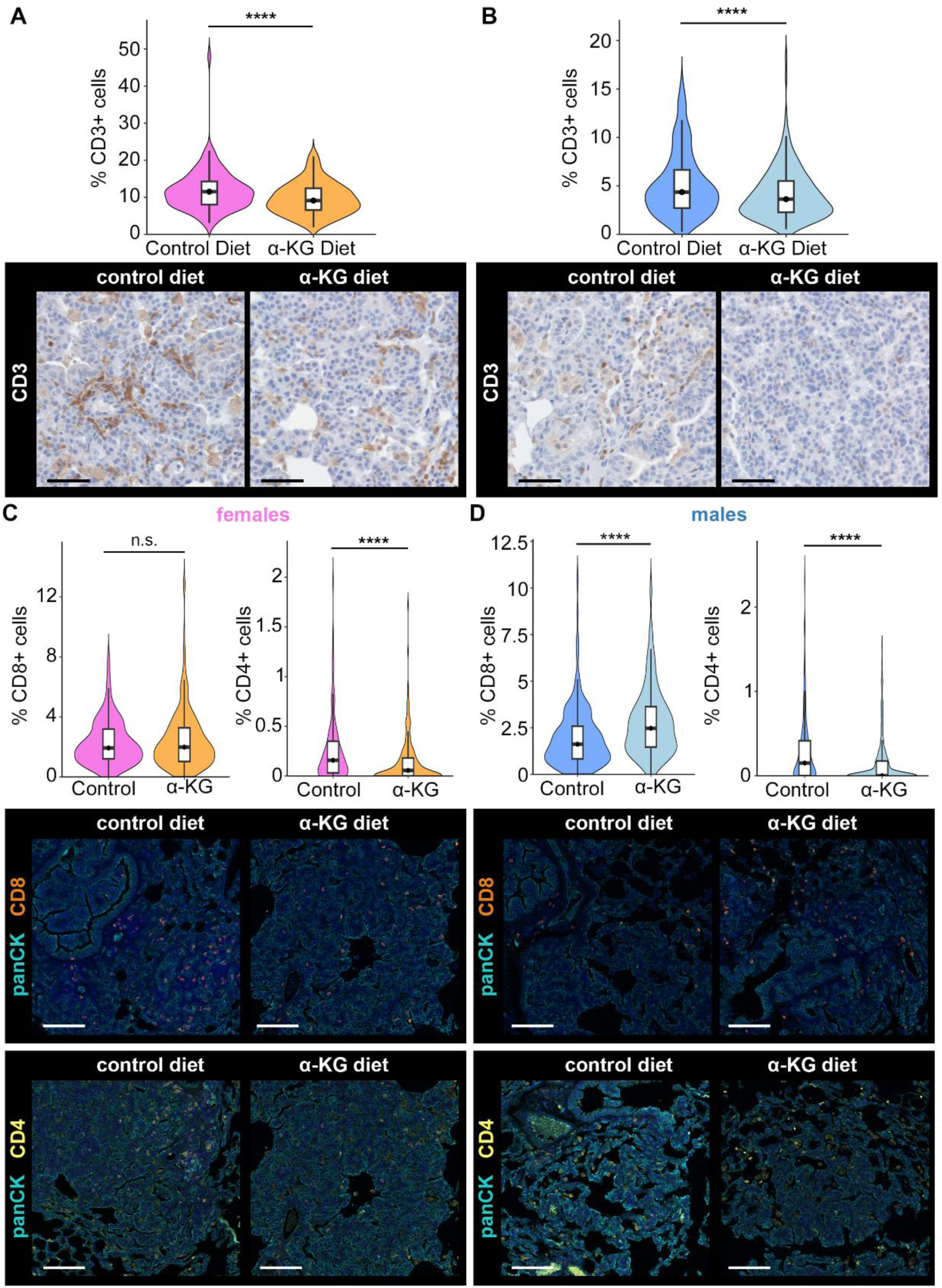
α-Ketoglutarate Remodels the Tumor Immune Microenvironment in a Sex-Dependent Manner. **(A-B)** Quantification of CD3^+^ T cells in female (A) and in male (B) LUAD tumors. Representative IHC images are reported at the bottom. **(C-D)** Multiplex immunofluorescence (mIF) analysis of T cell subsets: CD8^+^ and CD4^+^ T cells in female (C) and male (D) LUAD tumors. Representative mIF images of CD8+ T cells (orange), CD4+ T cells (yellow), and PanCK+ tumor cells (magenta) in female (C) and male (D) LUAD tumors. Data are shown as violin plots with overlaid box plots (median and interquartile range). Representative mIF images are from n = 4 mice per group. Scale bars for IHC = 50 µm. Scale bars for mIF = 100 µm.

To complement histological analyses, we inferred immune infiltration from bulk RNA-seq. Using published immune gene panels ^23,24^, we computed immune tumor microenvironment signature scores across major lymphoid and myeloid compartments. These revealed sex-dependent remodeling (Fig. S4B). In females, α-KG broadly reduced dendritic cells, macrophages, T cell exhaustion, and regulatory T cell (Treg) programs. In males, α-KG broadly increased macrophage and T cell exhaustion. This analysis suggests that α-KG may opposite effects on myeloid and T cell programs depending on sex.

### Wild-Type p53 Is Required for the Anti-Tumor Effect of Dietary α-Ketoglutarate

The anti-cancer effects of α-KG have previously been linked to p53 activity^17^. To test whether p53 is required for the observed reduction in tumor burden, we performed a treatment trial in KP mice (*Kras^LSL-G12D/+^*; *Trp53^flox/flox^*) on C57BL/6J background. In KP mice on dietary α-KG, females exhibited a non-significant trend toward increased tumor area (p = 0.0686), whereas males showed no effect (p = 0.4342). There were no significant differences in tumor burden at baseline between the two sexes (p = 0.2788) (Fig. S6).

These results suggest that α-KG requires wild-type p53 to exert its anti-cancer effects in LUAD. Loss of *Trp53* reversed the protective trend observed in female *Kras*-only models, consistent with prior work demonstrating that α-KG can act downstream of p53 to promote differentiation and tumor suppression^17^.

## DISCUSSION

To our knowledge, this is the first study to describe sex-dependent effects of α-Ketoglutarate (α-KG) supplementation on histone methylation, transcriptional programming, and immune contexture in LUAD. In a *KRAS*-driven mouse model, long-term α-KG supplementation was associated with reduced tumor area and proliferation in females but increased tumor area and proliferation in males. These findings suggest that sex is an important determinant of tumor responses to metabolic interventions.

At the histone level, α-KG was associated with reduced global levels of the repressive modifications H3K27me3 and H3K9me3 in females, consistent with a state more permissive to lineage-associated transcription^25–27^. In males, α-KG was instead associated with increased H3K27me3 without changes in H3K4me3 or H3K9me3, aligning with enhanced repression. As a cofactor for Jumonji-domain histone demethylases, α-KG may contribute to sex-dependent variation in demethylase activity^28–30^. Although not directly tested here, KDM6A represents one candidate mechanism that could contribute to the observed sex differences in histone methylation^31–33^. We previously showed that hypermethylation of H3K27, target of the KDM6A demethylase activity, is associated with a more aggressive phenotype in LUAD^22^. Lower KDM6A activity in males than in females could limit α-KG-driven demethylation, contributing to the accumulation of repressive marks. Testing this mechanism will be an important direction for future work.

In females, α-KG was associated with enrichment of a myogenesis signature, including genes such as *Myom2*, *Tnnt1*, and *Fabp3*, which are typically restricted to cardiac or skeletal muscle and rarely expressed in epithelial tumors^18–20^. In contrast, males exhibited higher baseline expression of the myogenesis signature, and α-KG treatment was associated with repression of the myogenesis genes. This opposing regulation suggests a sexually dimorphic transcriptional response, with α-KG promoting myogenesis-associated programs in females while suppressing them in males. Transcription factor enrichment analysis identified TBX5, a T-box family member involved in cardiac development and epithelial identity^21,34,35^, as a top candidate regulator. α-KG-treated female tumors showed increased TBX5 transcript and nuclear protein, concordant with myogenesis-related signatures. In contrast, nuclear TBX5 was reduced in male tumors, while nuclear GATA4 was selectively induced, suggesting engagement of alternative transcriptional circuits that may oppose lineage programs or reinforce tumor plasticity^36,37^. The TCGA data and the pattern of regulation by α-KG in mice support a model in which TBX5 marks a less proliferative LUAD state. Future studies will be needed to directly test whether TBX5 is required for the differentiation phenotype we observed.

Baseline analyses revealed that TBX5 was intrinsically higher in male tumors, whereas GATA4 was higher in female tumors. Prior studies demonstrate that TBX5 and GATA4 physically interact and co-occupy enhancers with other cardiac transcription factors to drive lineage-specific gene expression ^38^. Mutational and genetic evidence further supports dosage-sensitive interactions between these factors, where imbalanced levels can alter cardiac development and function ^39,40^. Moreover, TBX5 has been shown to exert dosage-sensitive, rheostatic control of cardiac gene expression, with graded levels dictating distinct morphogenetic outcomes ^41^. Consistent with this, TBX5 overexpression in non-small cell lung cancer cell lines suppresses proliferation and induces apoptosis^42^, supporting a functional role for TBX5 in restraining tumor growth.

α-KG was also associated with remodeling of the tumor immune microenvironment in a sex-dependent manner. Immunohistochemistry showed that overall CD3^+^ T cell infiltration was reduced in both sexes, while multiplex immunofluorescence revealed expansion of CD8^+^ T cells specifically in males. The expansion of CD8^+^ T cells in males was unexpected, as it coincided with increased tumor burden, a pattern described in other cancers where T cells expand but remain exhausted or dysfunctional ^43^. The transcriptional down-regulation of checkpoint genes *Pdcd1*, *Ctla4*, *Lag3* in females and upregulation of checkpoint gene *Tigit* in males supports this possibility ^44^, although functional assays will be required to determine whether these CD8^+^ cells are exhausted or clonally amplified. Consistent with this, immune signature analysis from bulk RNA-seq revealed that α-KG treatment increased T cell exhaustion programs in male tumors, supporting the possibility that the expanded CD8^+^ population is functionally impaired. A limitation of our study is that immune remodeling was primarily inferred from bulk RNA-seq and histological analyses; future work using single-cell or functional immunological assays will be necessary to directly establish the impact of α-KG on immune function.

At this time, we cannot explain why male mice treated with α-KG developed larger tumors than those on control diet. We believe that this finding is not generalizable because we performed the same trials in FVB/N *Kras^LSL-G12D/+^* mice, and in this genetic background α-KG did not change tumor burden in males (p = 0.8443), while still trending toward tumor reduction in females (p = 0.1096) (Fig. S6). This observation suggests that the worse tumor burden in C57BL/6 males may be due to a peculiarity of this genetic background.

These observations are consistent with established roles of sex chromosomes, hormone signaling, and tumor-immune crosstalk in shaping T cell activation and checkpoint expression ^45–49^, highlighting the importance of incorporating sex and tumor genotype as biological variables in metabolic-epigenetic cancer interventions. Future studies integrating chromatin state, transcriptional programs, and immune features will be essential to identify patient subsets most likely to benefit from α-KG-based approaches. Collectively, our findings position dietary α-KG as a candidate for metabolic-epigenetic intervention in lung cancer and highlight that such strategies must be deployed within precision frameworks that explicitly account for sex and genetic context. More broadly, our study points to a novel principle: metabolites such as α-KG can drive divergent chromatin and immune outcomes depending on biological context, underscoring the need for precision frameworks that account for sex and tumor genotype in their evaluation.

## STAR★METHODS

### EXPERIMENTAL MODEL AND STUDY PARTICIPANT DETAILS

Mice carrying a conditional *Kras^LSL-G12D/+^* allele on a C57BL/6J background were obtained from Jackson Laboratories. *Kras^LSL-G12D/+^*; *Trp53^flox/flox^* (KP) mice were bred in out colony from *Kras^LSL-G12D/+^* and *Trp53^flox/flox^* strains also obtained from Jackson Laboratories. *Kras^LSL-G12D/+^* mice on an FVB/N background were kindly provided by Dr. David Shackelford. At twelve weeks of age, both male and female mice were anesthetized with ketamine/xylazine and administered Ad5-CMV-Cre virus intranasally (1 × 10^7 PFU/µL, 15 µL per nostril) to initiate lung tumorigenesis. One week post-infection, mice were randomized by age and sex into experimental groups and placed on either a control diet (Envigo #7013) or a diet with the same composition but supplemented with 2% (w/w) calcium α-ketoglutarate (Ca-αKG) (Carbosynth #71686-01-6; Envigo TD.230157). The B6 *Kras* and FVB *Kras* cohorts were maintained on their assigned diets for 16 weeks, whereas KP mice were maintained for 12 weeks. All animal procedures were approved by the UCLA Animal Research Committee and conducted in compliance with institutional guidelines.

### METHOD DETAILS

#### Histology and Immunohistochemistry

Lungs were inflated with 10% neutral buffered formalin, fixed overnight at room temperature, dehydrated in graded ethanol, cleared in xylene, and paraffin embedded. Sections were cut at 5 µm, deparaffinized, and rehydrated. For H&E staining, sections were stained with hematoxylin and counterstained with eosin before mounting.

For immunohistochemistry, antigen retrieval was performed using citrate buffer (AR6, pH 6) or Tris-EDTA buffer (AR9, pH 9). Primary antibodies included TBX5 (Sino Biological #107330-T08, 1:10000, AR9), GATA4 (ab307823, 1:100, AR9), and CD3 (Dako A0452, 1:2000, AR6). Endogenous peroxidase activity was quenched using hydrogen peroxide, and detection was performed with the ImmPRESS® Horse Anti-Rabbit IgG PLUS Polymer Kit (Vector Labs MP-7401). Slides were developed with DAB chromogen, counterstained with Harris hematoxylin, dehydrated, and mounted. Whole-slide images were acquired on the Vectra Polaris imaging system, and quantification of immune infiltration and H-scores was performed using QuPath (v0.4.3) with custom pixel classifiers and nuclear detection parameters.

#### Multiplex Immunofluorescence

Multiplex immunofluorescence (mIF) was performed using the Ventana Discovery Ultra (Roche) and Opal fluorophores (Akoya Biosciences). Five-micrometer tissue sections on Superfrost slides were deparaffinized with EZ-Prep reagent (Roche) and subjected to antigen retrieval in CC1 buffer (pH 9, 95 °C). Discovery Inhibitor (Roche) was applied, followed by sequential rounds of staining with CD4, CD8, panCK, and Ki67. Each round consisted of a primary antibody incubation followed by OmniMap secondary antibody detection (Roche). Signal amplification was achieved with Opal fluorophores (570, 620, 650, 480). Between rounds, sections underwent heat-induced epitope retrieval to strip prior antibody complexes. Slides were counterstained with Spectral DAPI and mounted in ProLong Diamond antifade medium (Thermo Fisher). Imaging was performed using the Vectra Polaris. Images were spectrally unmixed with inForm (Akoya), stitched and analyzed in HALO (Indica Labs), and subjected to whole-slide spatial analysis. Cell phenotyping and quantification were performed with a uniform threshold across all slides. All quantifications were blinded to treatment group and sex, and data were exported and graphed in R.

#### Tumor Burden Quantification

Tumor burden was quantified from H&E-stained lung sections. Whole-slide images were obtained with the Vectra Polaris and analyzed using QuPath. Tumor regions were manually annotated using the magic wand tool based on histological features, and tumor area was expressed as the percentage of total lung area occupied by tumor. All quantifications were performed blinded to treatment and sex.

#### RNA Sequencing and Analysis

RNA-seq libraries were prepared using the KAPA Stranded mRNA-Seq Kit. The workflow included mRNA enrichment and fragmentation, cDNA synthesis with random priming, second-strand synthesis incorporating dUTP, end repair, A-tailing, adaptor ligation, and PCR amplification. Libraries were sequenced on an Illumina NovaSeq X Plus with a paired-end 2 × 50 bp configuration. Data quality was assessed using Illumina SAV, and demultiplexing was performed with bcl2fastq v2.19.1.403. Alignment was carried out using STAR aligner (v2.7) with the GRCm39 reference genome and Ensembl release 108 annotation. Transcript counts were normalized using the median ratio method in Partek Flow. Differential expression was determined with DESeq2. Pathway enrichment of significant genes was performed using ShinyGO v0.8. Gene set enrichment analysis (GSEA) was performed with the MSigDB Hallmark collection, and transcription factor enrichment was performed using ChEA3. Full differential expression results and transcription factor enrichment output are provided in Supplementary Tables S1 and S2.

#### Western Blotting for Histone Modifications

Histones were extracted from snap-frozen tumor tissue using the Abcam Histone Extraction Kit (ab113476). Protein concentration was measured using the BCA assay. Twenty micrograms of histone protein per sample were resolved on NuPAGE Bis-Tris 4-12% gels and transferred to PVDF membranes. Blots were probed with primary antibodies against H3K27me3, H3K4me3, and H3K9me3 (CST, 1:2000 each). Chemiluminescent detection was used, and band intensities were quantified in ImageJ. Membranes were stained with Coomassie Brilliant Blue to control for total histone content, and band intensities were normalized accordingly.

#### TCGA LUAD Analysis

Publicly available TCGA-LUAD data were accessed through UCSC Xena. TBX5 expression and selected gene sets were stratified by sex. Survival analyses were conducted using median-split expression groups. Kaplan-Meier survival curves and Cox proportional hazards models were performed in R using the survival package. Boxplots, scatterplots, and Spearman correlations were generated with ggplot2.

### QUANTIFICATION AND STATISTICAL ANALYSIS

All statistical analyses for mouse experiments were performed in R (v4.3.2). Normality was assessed using the Shapiro-Wilk test. For normally distributed data, unpaired t-tests were used; for non-normal distributions, Wilcoxon rank-sum tests were applied. Significance was defined as p < 0.05.

For human data, analyses were conducted using TCGA LUAD datasets. Kaplan–Meier survival curves, Cox proportional hazards models, and correlation analyses were generated in R using the survival and ggplot2 packages. All analyses were stratified by sex. Quantifications from imaging experiments were performed blinded to sex and treatment group.

### ADDITIONAL RESOURCES

This study did not generate additional resources.

## Resource Availability

### Lead contact

Further information and requests for resources and reagents should be directed to and will be fulfilled by the lead contact, Claudio Scafoglio.

### Materials availability

This study did not generate new unique reagents.

### Data and code availability

The RNA-seq raw data are publicly available in ArrayExpress repository under accession number:

E-MTAB-15597.

Any additional information required to re-analyze the data reported in this paper is available from the lead contact upon request.

## Acknowledgements and Funding

RNA sequencing and bioinformatic analysis were conducted with support from the UCLA Technology Center for Genomics & Bioinformatics (TCGB). Histological processing and slide preparation were performed by the UCLA Translational Pathology Core Laboratory (TPCL). We thank Dr. David Shackelford (UCLA) for providing the *Kras*-mutant mice in FVB background. We thank Dr. Orian Shirihai for critical advice and mentorship. Figures were prepared with the aid of Biorender (https://BioRender.com).

This work was supported in part by the NIH/NCI R01 grant R01CA237401-01A1, the American Cancer Society Discovery Boost Grant DBG-23-1152703-01-TBE, and the LUNGevity Foundation Lung Cancer Early Detection Award #857709 (to C.S.). M.A. is supported by the UCLA Tumor Immunology Training Grant (Ruth L. Kirschstein Institutional NRSA T32 CA009120), the Gates Millennium Scholars Program through the Bill and Melinda Gates Foundation, and the Eugene V. Cota-Robles Fellowship at UCLA. M.A. also acknowledges the support of his family and community - especially those in East Los Angeles - for inspiring his path in science.

## Author Contributions

**Conceptualization:** M.A.J., C.S.

**Methodology:** M.A.J., C.S.

**Investigation:** M.A.J., M.S., L.P., E.G., A.S., C.D., A.P., P.S.

**Formal Analysis:** M.A.J., C.S.

**Data Curation:** M.A.J., C.S.

**Writing - Original Draft:** M.A.J.

**Writing - Review & Editing:** M.A.J., C.S., B.L.

**Supervision:** C.S.

**Resources:** C.S., B.L., S.M.D.

**Funding Acquisition:** M.A.J., S.M.D., C.S.

## Declaration of interests

The authors declare no competing interests.

## Declaration of generative AI and AI-assisted technologies

The authors declare no use of generative AI and AI-assisted technologies.

## Supplemental Information

Document S1: Figures S1-S6

Supplementary Tables: Tables S1-S2

---

**Figure S1.**
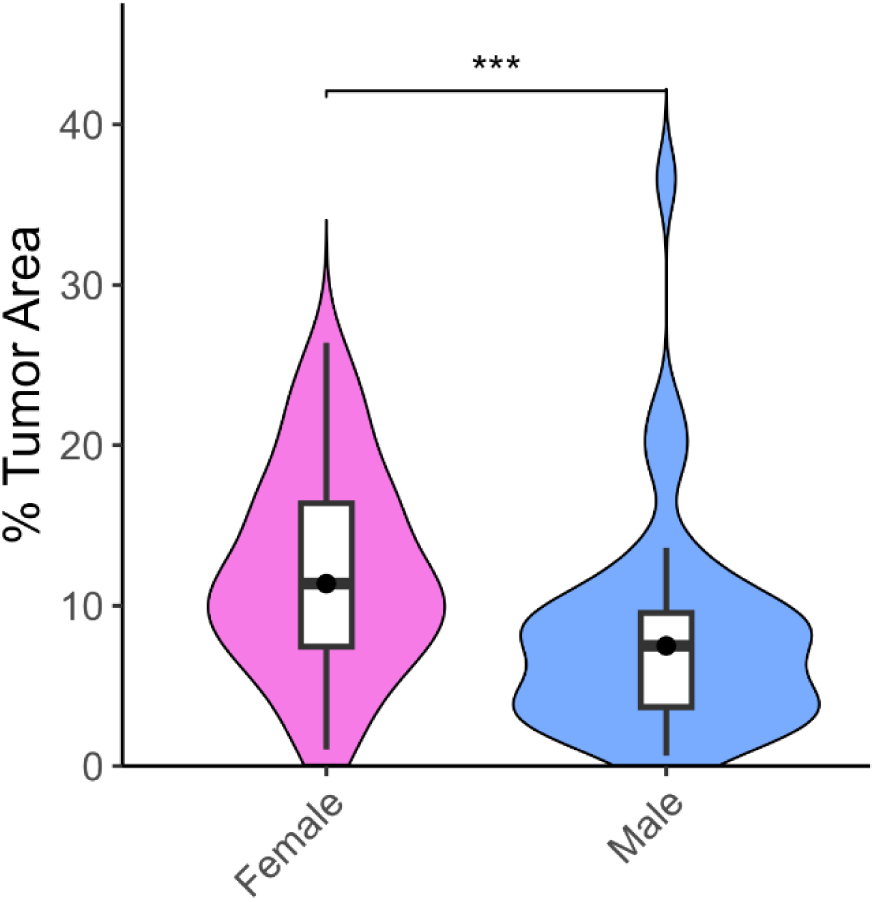
In the Kras*^G12D^*-driven model (C57BL/6 background) Female LUAD Tumors are Significantly Larger than Male Tumors at Baseline. Tumor burden was quantified in LUAD-bearing mice on control diet at 17 weeks post-AdCre induction. Violin plots show percentage of tumor area in female (pink) and male (blue) mice. Significance was measured by Wilcoxon rank-sum test.

**Figure S2.**
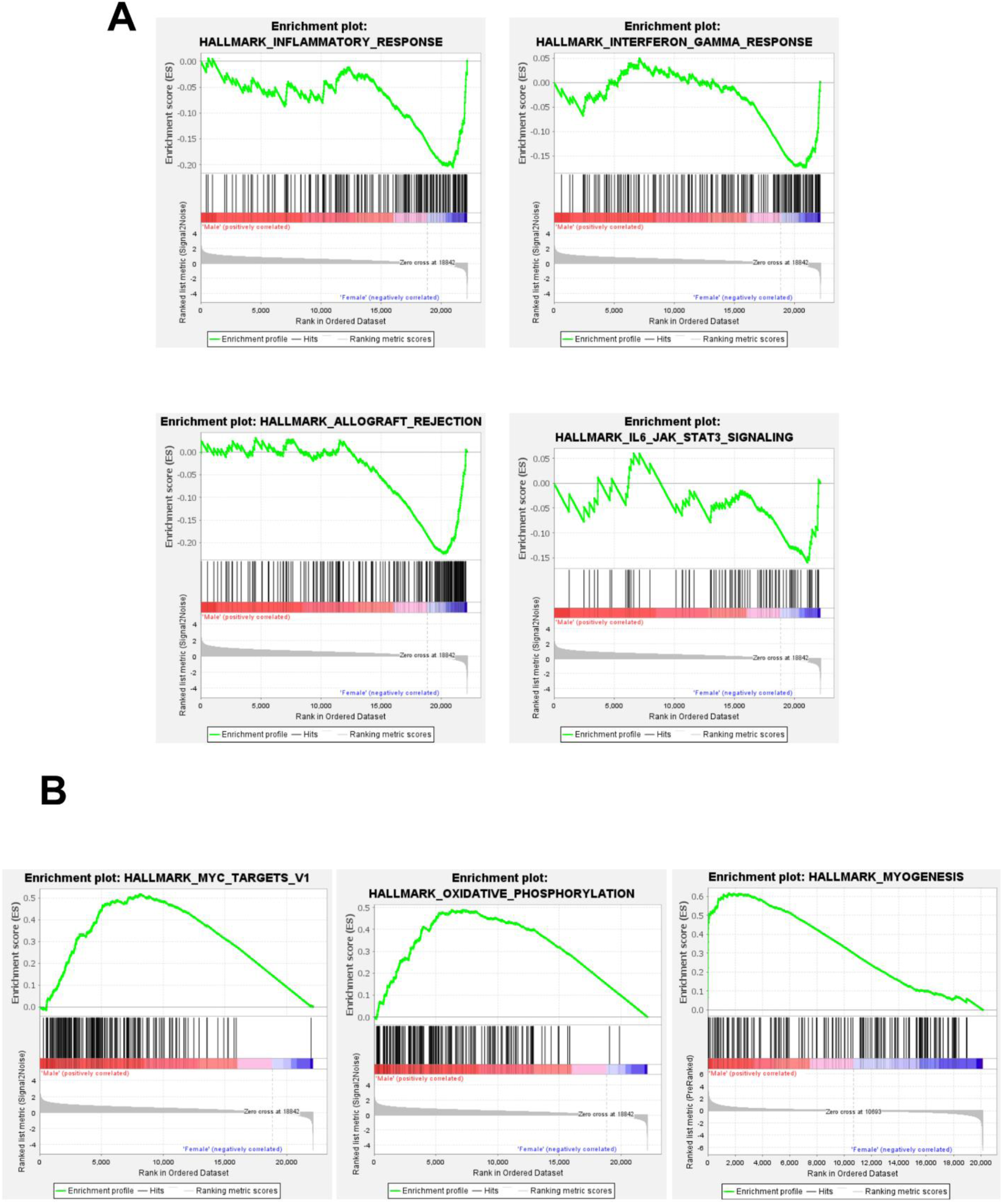
Basal Sex Differences in LUAD Tumors Highlight Immune Suppression in Females and Metabolic Activation in Males. **(A)** Gene sets enriched in female tumors: INFLAMMATORY_RESPONSE (A, NES = –0.79, q < 0.001), INTERFERON_GAMMA_RESPONSE (NES = –0.72, q < 0.001), ALLOGRAFT_REJECTION (NES = –0.76, q < 0.001), and IL6_JAK_STAT3 (NES = –0.72, q = 0.002). **(B)** Gene sets enriched in male tumors: MYC (NES = 1.41, q < 0.001), OXIDATIVE_PHOSPHORYLATION (NES = 1.46, q < 0.001), and MYOGENESIS (NES = 2.3, q < 0.001). Data are from bulk RNA sequencing of lung tumors harvested at 17 weeks (n = 4 tumors per group). Differential expression was performed using DESeq2. GSEA was performed using the Hallmark gene set collection (MSigDB v7.5.1). NES = normalized enrichment score.

**Figure S3.**
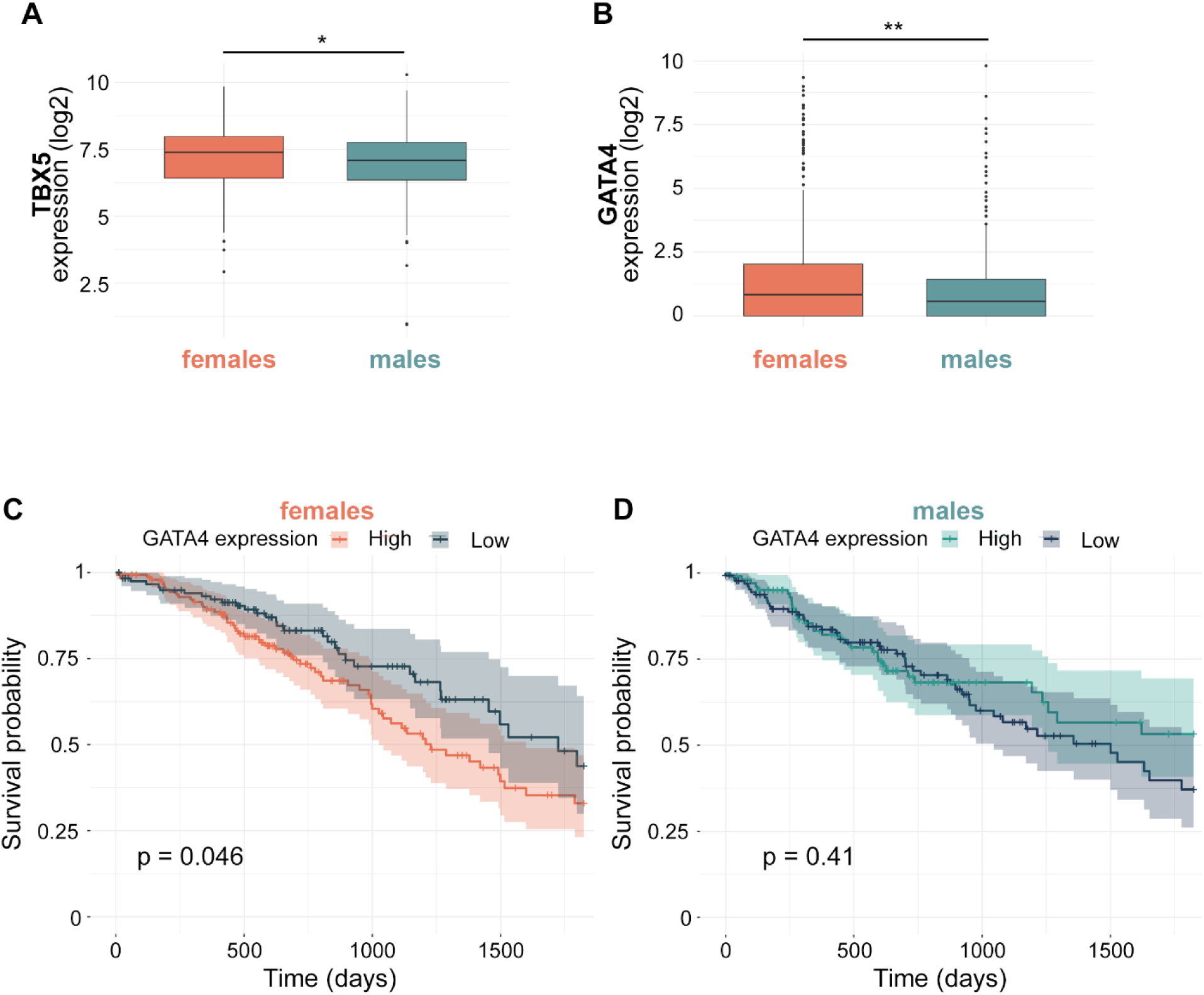
Female-Biased GATA4 Expression in LUAD Predicts Poorer Survival in Females but Not in Males. **(A)** TBX5 expression in female and male LUAD tumors from the TCGA dataset (Wilcoxon rank-sum test, *p = 0.035*). **(B)** GATA4 expression in female and male LUAD tumors from the TCGA dataset (Wilcoxon rank-sum test, *p = 0.0025*). **(C)** Kaplan–Meier 5-year survival curves stratified by GATA4 expression (median split) in female LUAD patients (log-rank *p = 0.046*). **(D)** Kaplan–Meier 5-year survival curves stratified by GATA4 expression (median split) in male LUAD patients (log-rank *p = 0.41*).

**Figure S4.**
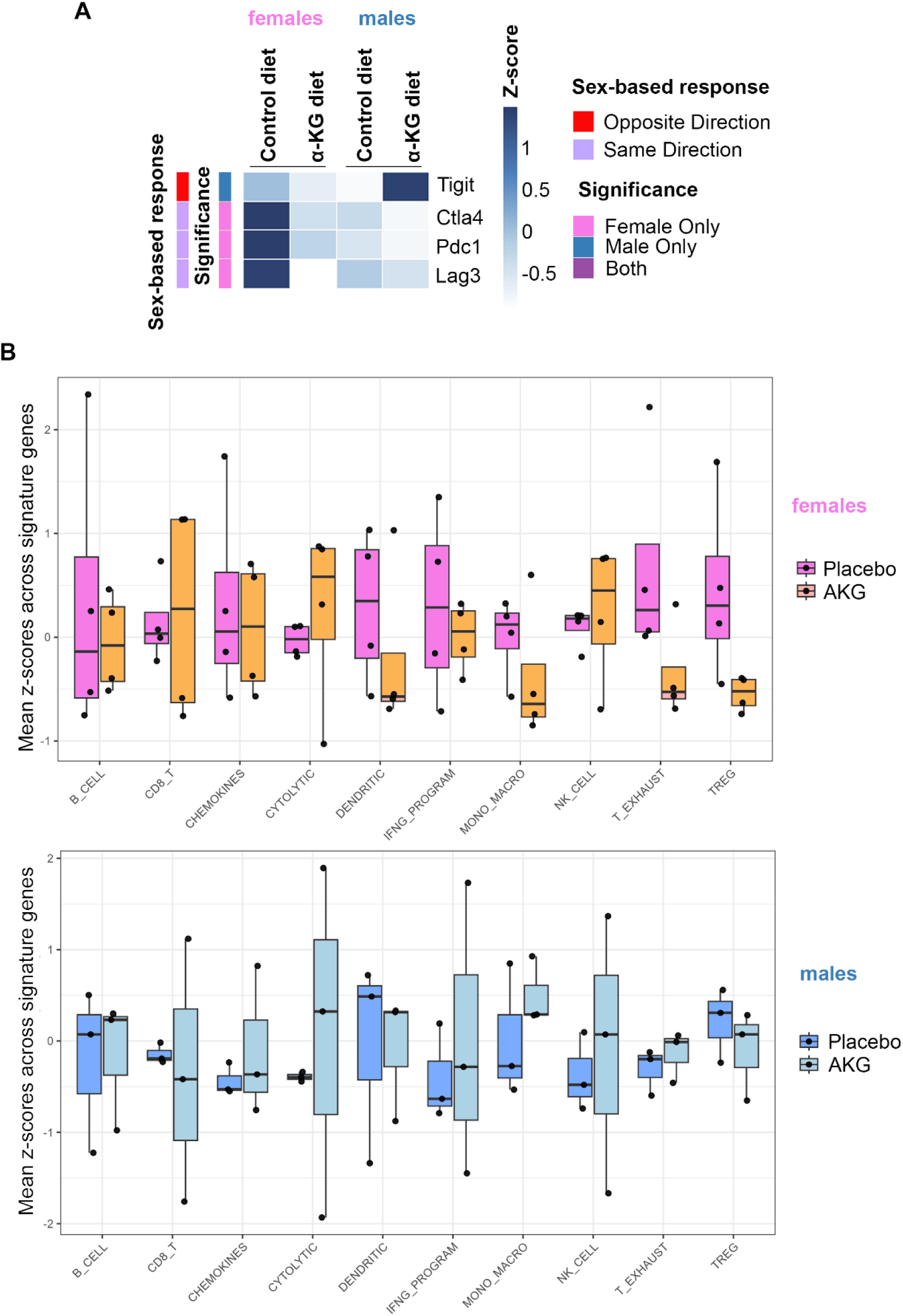
Immune Tumor Microenvironment (TME) Signature Scores from Bulk RNA-seq Reveal Sex-Dependent Remodeling by α-KG. **(A)** Heatmap of immune chackpoint-related transcript expression from bulk RNA-seq. **(B)** Immune TME signature scores derived from published gene panels were calculated from LUAD bulk RNA-seq data as the mean z-score across signature genes. Boxplots show sex-stratified comparisons between normal diet and α-KG diet. Upper panel: female LUAD tumors (n = 4 per group). Lower panel: male LUAD tumors (n = 3 per group). Signatures include B cells, CD8⁺ T cells, chemokines, cytolytic program, dendritic cells, IFN-γ program, macrophages, NK cells, T cell exhaustion, and Tregs. Data are shown as boxplots with individual sample points (median and interquartile range).

**Figure S5.**
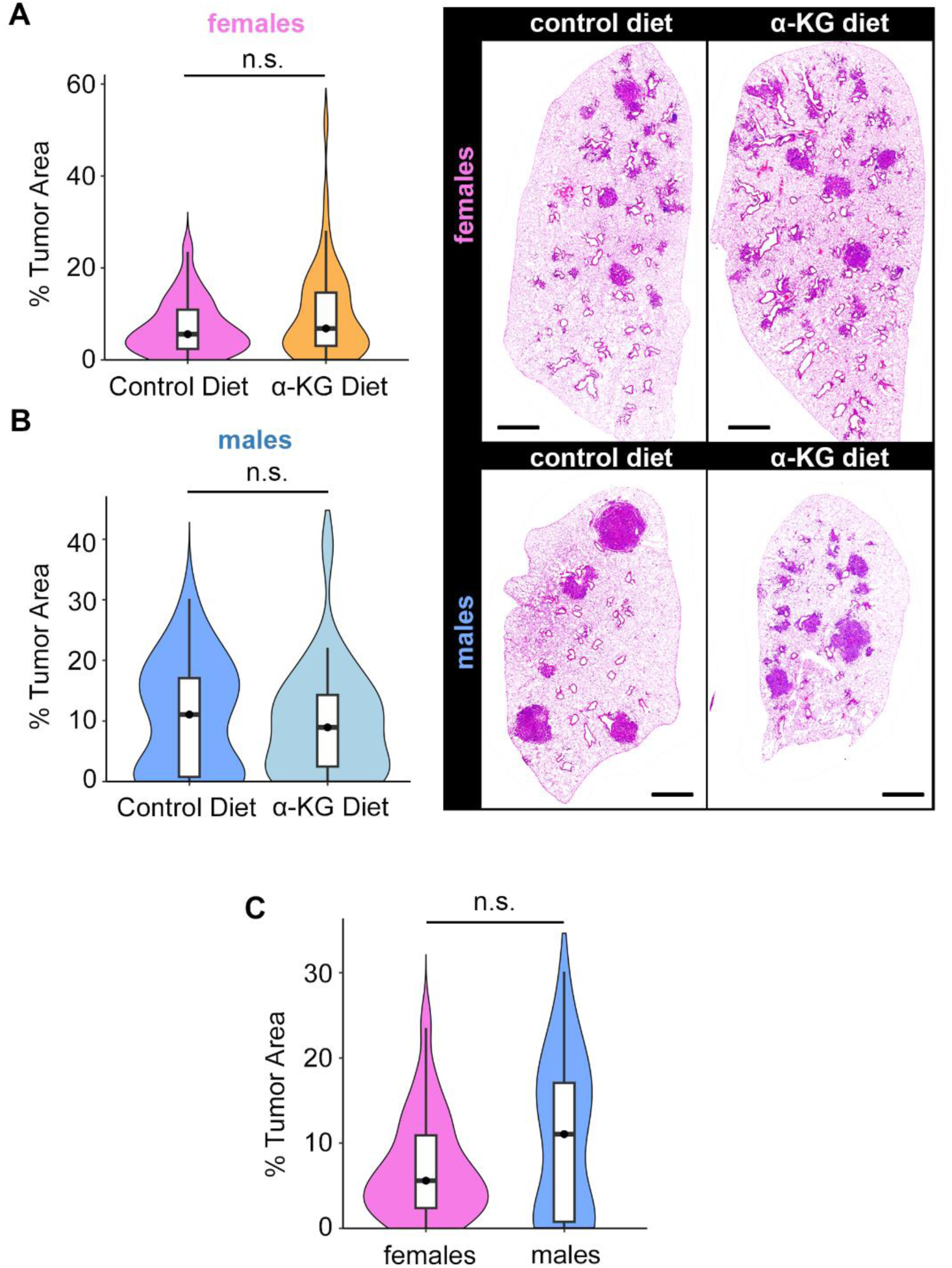
Wild-Type p53 Is Required for the Anti-Tumor Effect of Dietary α-Ketoglutarate. **(A)** Quantification of tumor area per lobe in female KP mice on control diet versus Ca-αKG diet (unpaired *t*-test; *p* = 0.06857; n = 11 mice for control diet, 10 mice for α-KG). **(B)** Quantification of tumor area per lobe in male KP mice on control diet versus Ca-αKG diet (Wilcoxon rank-sum test; *p* = 0.4342; n = 6 mice for control diet, 5 mice for α-KG). **(C)** Comparison of tumor burden between female and male KP on control diet (Wilcoxon rank-sum test; *p* = 0.2788). Data are presented as violin plots with overlaid box plots (median and interquartile range). Tumor area was quantified as the percentage of lung tissue occupied by tumors manual annotation.

**Figure S6.**
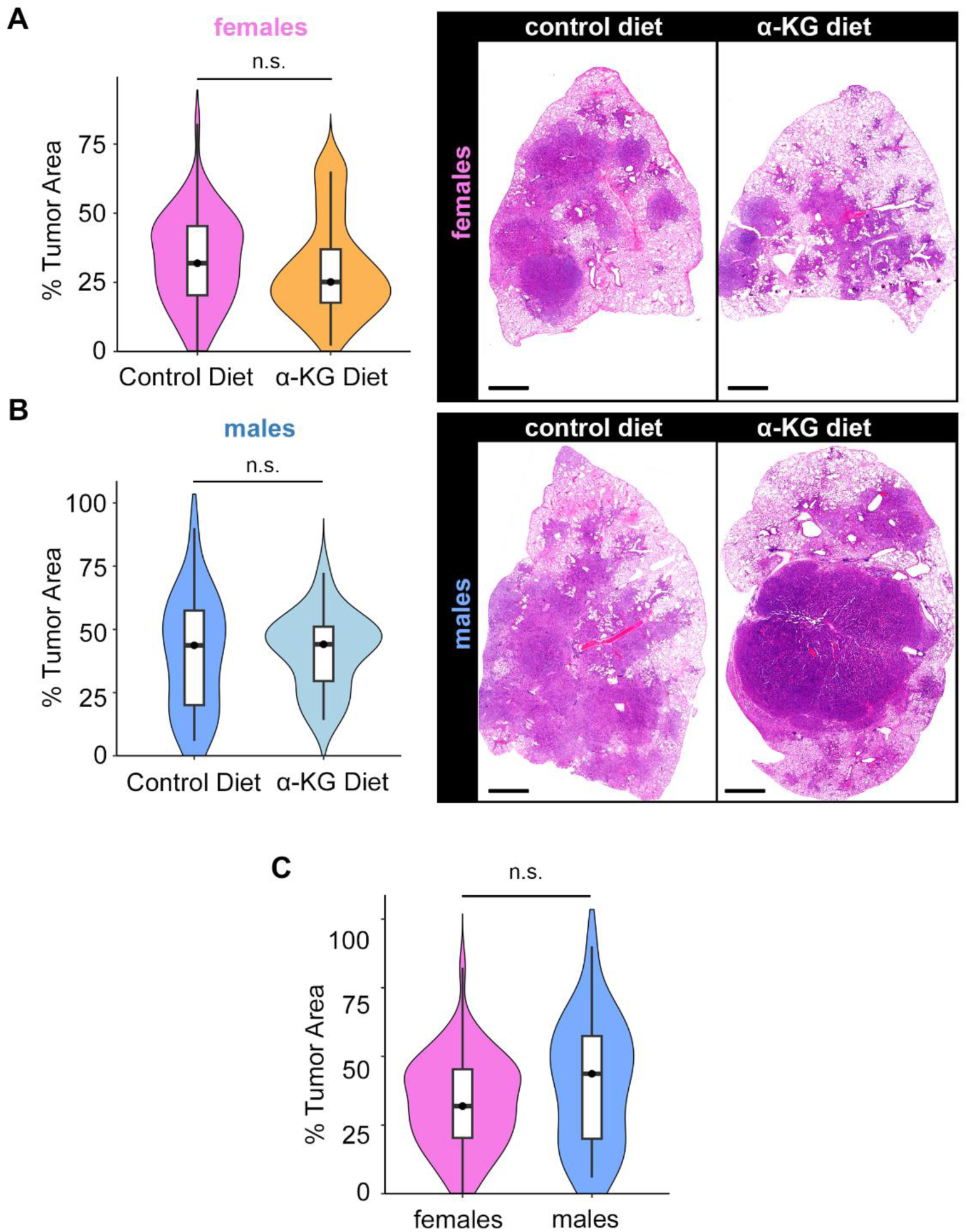
Dietary α-Ketoglutarate does not significantly alter tumor burden in FVB/N Kras mice. **(A)** Quantification of tumor area per lobe in female FVB/N Kras mice on control diet versus Ca-αKG diet (unpaired *t*-test; *p* = 0.1096; n = 16 mice for control diet, 17 mice for α-KG). **(B)** Quantification of tumor area per lobe in male FVB/N Kras mice on control diet versus Ca-αKG diet (unpaired *t*-test; *p* = 0.8443; n = 5 mice for control diet, 5 mice for α-KG). **(C)** Comparison of tumor burden between female and male FVB/N Kras mice on control diet (unpaired *t*-test; *p* = 0.1194). Data are presented as violin plots with overlaid box plots (median and interquartile range). Tumor area was quantified as the percentage of lung tissue occupied by tumors manual annotation.

## Notes

### Competing Interest Statement

The authors have declared no competing interest.

